# A 6-CpG Validated Methylation Risk Score Model for Metabolic Syndrome: The HyperGEN and GOLDN Studies

**DOI:** 10.1101/2021.10.22.465467

**Authors:** Bertha A Hidalgo, Bre Minniefield, Amit Patki, Rikki Tanner, Minoo Bhagheri, Hemant K. Tiwari, Donna K. Arnett, M. Ryan Irvin

## Abstract

There has been great interest in genetic risk prediction using risk scores in recent years, however, the utility of scores developed in European populations and later applied to non-European populations has not been successful. In this study, we used cross-sectional data from the Hypertension Genetic Epidemiology Network (HyperGEN, N=614 African Americans (AA)) and the Genetics of Lipid Lowering Drugs and Diet Network (GOLDN, N=995 European Americans (EA)), to create a methylation risk score (MRS) for metabolic syndrome (MetS), demonstrating the utility of MRS across race groups. To demonstrate this, we first selected cytosine-guanine dinucleotides (CpG) sites measured on Illumina Methyl450 arrays previously reported to be significantly associated with MetS and/or component conditions (*CPT1A* cg00574958, *PHOSPHO1* cg02650017, *ABCG1* cg06500161, *SREBF1* cg11024682, *SOCS3* cg18181703, *TXNIP* cg19693031). Second, we calculated the parameter estimates for the 6 CpGs in the HyperGEN data and used the beta estimates as weights to construct a MRS in HyperGEN, which was validated in GOLDN. We performed association analyses using a logistic mixed model to test the association between the MRS and MetS adjusting for covariates. Results showed the MRS was significantly associated with MetS in both populations. In summary, a MRS for MetS was a strong predictor for the condition across two ethnic groups suggesting MRS may be useful to examine metabolic disease risk or related complications across ethnic groups.

## Introduction

Genome wide association studies (GWAS) have examined the cumulative effect of novel, single variants on both quantitative trait variance and disease status, by summing up the number of risk alleles at each locus weighted by the effect size at each locus, otherwise known as a genetic risk score (GRS) [1]. While once limited by small sample sizes, GRSs have since benefitted from the growth and development of large-disease based consortia, along with improved methodology which allow for the aggregation of thousands to millions of genetic variants (i.e. polygenic risk scores, PRS), which better inform risk prediction [2]. A major limitation of GRS – including PRS - is that they have been developed and optimized for European-ancestry populations, thus limiting their utility and generalizability in non-European ancestry populations. Recognizing these limitations, we aimed to apply statistical approaches common to GRS and PRS to epigenome-wide association (EWAS) data to evaluate if MRS may further enhance accuracy for complex disease prediction. To do so, we hypothesized that leveraging existing EWAS data and on previously reported associations between cytosine-phosphate-guanine (CpG) sites and complex disease traits, like metabolic syndrome, can improve prediction capabilities. To date, despite impressive effect sizes, strong statistical significance, and successful external replication in the EWAS literature (even across race/ethnic groups), few studies have examined the polygenomic effects of CpG sites (e.g., methylation risk scores (MRS)) on complex diseases.

As such, we applied this statistical approach for the prediction of metabolic syndrome in a population of African ancestry individuals by constructing an MRS, optimizing it and then validating it in a European ancestry population. Metabolic syndrome (MetS) is a cluster of conditions that can increase risk for cardiometabolic diseases. Early identification of MetS can help prevent onset of cardiometabolic disease later in life. A growing body of research has identified a number of CpGs that are associated with the multiple components of MetS. In this study, we leverage previously reported CpGs that have been significantly associated with conditions comprising the metabolic syndrome and/or closely related metabolic traits: body mass index [3, 4], waist circumference [4], dyslipedemia [5], fasting blood glucose [6], systolic and diastolic blood pressure [7], and HDL cholesterol [8, 9], to construct the MRS. Independent CpGs included were cg00574958 from carnitine palmitoyl transferase 1A (*CPT1A)*, cg02650017 from Phosphoethanolamine/Phosphocholine Phosphatase 1 (PHOSPHO1), cg06500161 from ATP Binding Cassette Subfamily G Member 1 (ABCG1), cg11024682 from Sterol Regulatory Element Binding Transcription Factor 1 (SREBF1), cg18181703 from Suppressor Of Cytokine Signaling 3 (SOCS3), and cg19693031 from Thioredoxin Interacting Protein (TXNIP). We used a weighted sum method to create the score among African Americans from the Hypertension Genetic Epidemiology Network (HyperGEN), and validated the score in European Americans from Genetics of Lipid Lowering Drugs and Diet Network (GOLDN).

## Methods

### Discovery and Validation Study Populations

Data for the discovery phase of this study was obtained from the HyperGEN study. HyperGEN is a cross-sectional study including over 1900 African-Americans from families, which included at least two siblings with hypertension onset before age 60 [10]. The study purpose was to examine possible interactions between genetic and non-genetic determinants of hypertension. In 2015, an ancillary epigenetic study was conducted on stored HyperGEN samples in the upper and lower tertial of echocardiography measured left ventricular mass [11]. After excluding those missing relevant phenotype data as previously described [12] a total 614 participants were included in the analysis. Both HyperGEN and GOLDN studies were approved by the Institutional Review Board at the University of Alabama at Birmingham.

Validation was conducted in GOLDN study [13]. European ancestry families in GOLDN were recruited from the Family Heart Study at two centers, Minneapolis, MN and Salt Lake City, UT to participate in a diet and/or drug intervention. In each case, only families with at least two siblings were recruited and only participants who did not take lipid-lowering agents (pharmaceuticals or nutraceuticals) for at least 4 weeks prior to the initial visit were included. For the present study, 994 GOLDN participants had available methylation data for a validation study of the HyperGEN MRS. Sample characteristics as well as clinical and lifestyle factors were considered in HyperGEN and GOLDN, including blood pressure, antihypertensive and lipid lowering medications, fasting blood glucose, triglycerides, HDL cholesterol, height, weight, and waist circumference have been described [7-9]. We used the published joint harmonized criteria to define MetS in HyperGEN in both studies [14].

### DNA Methylation and Data processing

#### HyperGEN

The Illumina HumanMethylation450 array was used to analyze DNA extracted from buffy coat at > 480,000 cytosine-phosphate-guanine (CpG) sites. Briefly, 500 ng of buffy coat DNA was hybridized to the Methyl450 array after bisulfite conversion with EZ DNA kits (Zymo Research, Irvine, CA) prior to standard Illumina amplification, hybridization, and imaging steps. The resulting intensity files were analyzed with Illumina’s GenomeStudio, which generated beta (*β*) scores (i.e., the proportion of total signal from the methylation-specific probe or color channel) and “detection *p* values” (probability that the total intensity for a given probe falls within the background signal intensity). Quality control (QC) measures were conducted by removing samples having more than 1% of CpG sites with a detection *p* value > 0.05, removing CpG sites having more than 5% of samples with a detection *p* value > 0.01, and individual CpG sites with detection *p* value > 0.01 set as missing. After these QC filters, 484,366 CpG sites were eligible for analysis. We normalized the *β* scores using the Subset-quantile Within Array Normalization (SWAN) method in *minifi* package to correct for differences between batches and the type I and type II assay designs within a single 450K array [15]. Cell count proportions (CD8 T lymphocytes, CD4 T lymphocytes, natural killer cells, B cells, and monocytes) were created using the algorithm developed by Houseman, which predicts underlying cellular composition of each sample from DNA methylation patterns [16].

#### GOLDN

CD4+ T-cells were isolated from frozen buffy coat samples isolated from peripheral blood collected at the baseline visit (prior to intervention). DNA was extracted using DNeasy kits (Qiagen, Venlo, Netherlands). 500ng of each DNA sample was treated with sodium bisulfite (Zymo Research, Irvine, CA). Normalization was performed on random subsets of 10,000 CpGs per run, with each array of 12 samples used as a “batch.” Probes from the Infinium I and II chemistries were separately normalized and β scores for Infinium II probes were then adjusted using the equation derived from fitting a second order polynomial to the observed methylation values across all pairs of probes located <50bp apart (within-chemistry correlations >0.99), where one probe was Infinium I and one was Infinium II. The filtered β scores were normalized using the ComBat R-package.

#### CgG candidate selection

We identified candidate DNA methylation loci, which have been currently and previously identified by large-scale EWAS comparing various ethnicities and cardiometabolic clinical characteristics. Based on the literature we chose six robust CpGs to construct our MRS: cg00574958 within *CPT1A*, cg02650017 within *PHOSPHO1*, cg06500161 within *ABCG1*, cg11024682 within *SREBF1*, cg18181703 within *SOCS3*, and cg19693031 within *TXNIP*. See Table 1 for references for genes selected.

**Table 1.**
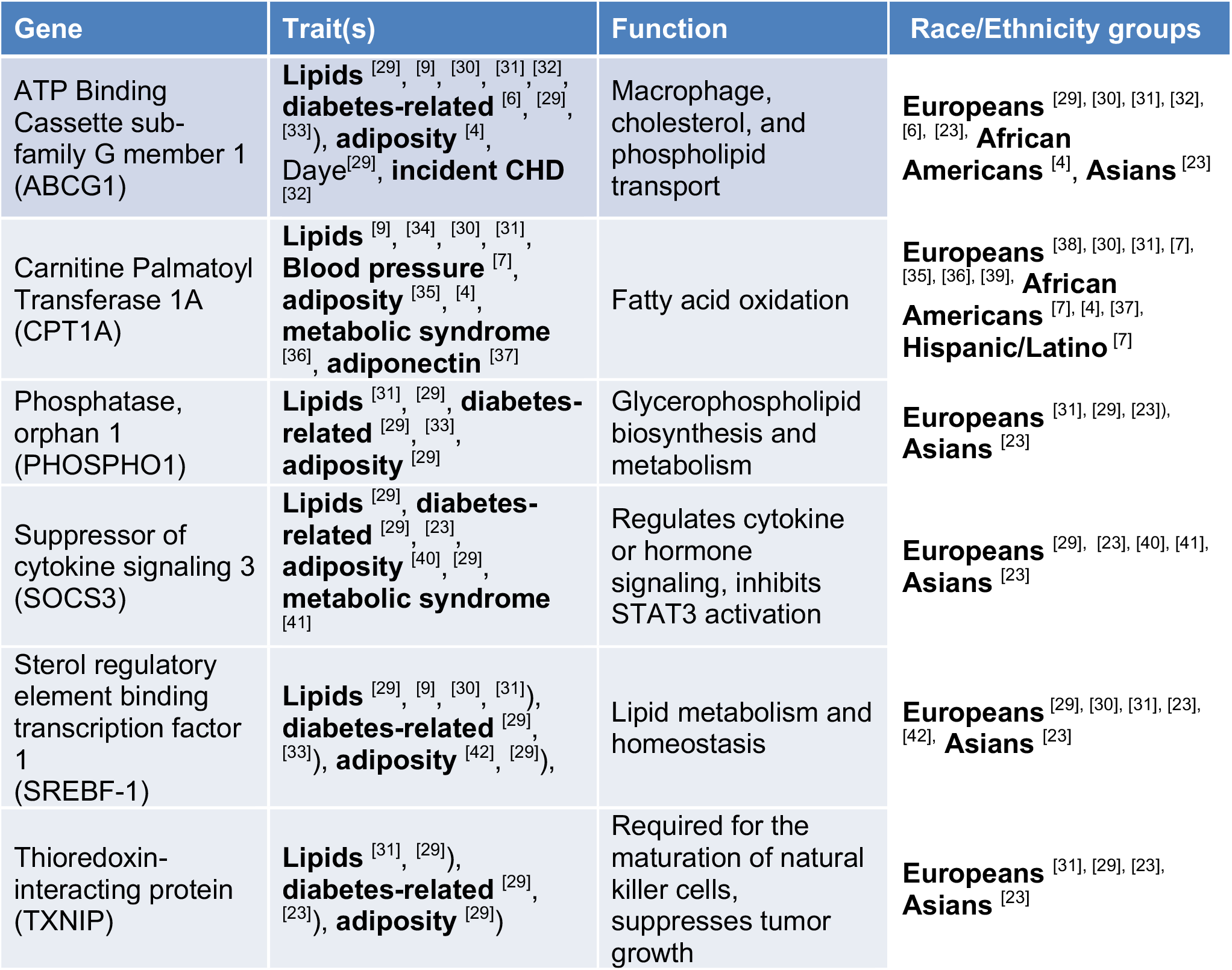
CPG Selection Literature Review.

#### Methylation Risk Score Model Building

A logistic mixed model was used to test the association between methylation at each candidate CpG site and MetS in HyperGEN. We adjusted for age, sex, study site, and estimated blood cell counts as fixed effects, and family structure as a random effect. Parallel models were implemented in GOLDN except methylation principal components replaced the estimated blood cell counts to adjust for cell type impurity (GOLDN was of a single cell type). To calculate the MRS, Z-values from the candidate CpG HyperGEN models described above (shown in Table 2) were utilized as weights and multiplied by the CpG-values and the product was summed to generate a risk score for each sample ((Z_1_* cpg_1 beta score_) + (Z_2_* cpg_2 beta score_)+ … (Z6* cpg_6 beta score_)). The mean and standard deviation of the score in HyperGEN was calculated. Each participants MRS was then standardized by subtracting the mean and dividing by the standard deviation. We used a parallel approach to calculate the standardized MRS in GOLDN by using GOLDN CpG values weighted by the HyperGEN Z-values.

**Table 2.**
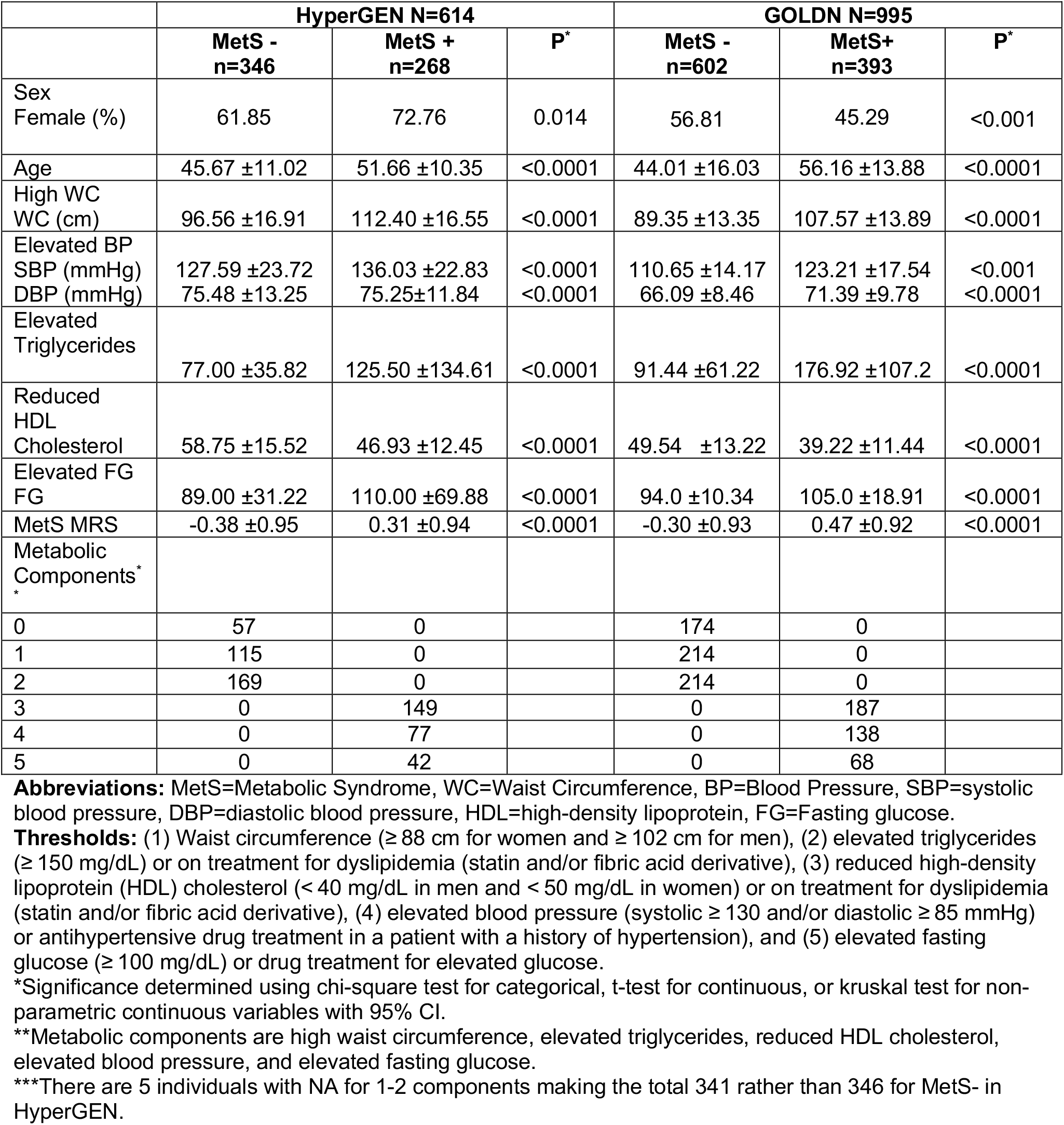
Baseline Characteristics of HyperGEN and GOLDN Study Participants.

### Methylation Risk Score Model Performance Testing

We compared both HyperGEN and GOLDN characteristics between individuals with (MetS+) and without (MetS-) metabolic syndrome. Significance of these characteristics were calculated using a simple t test for continuous traits and a chi-square test for binary traits. We then tested the association between the standardized MRS and MetS (outcome) using a logistic mixed model in HyperGEN adjusting for age, sex, study site, 4 ancestry principal components, estimated blood cell counts as fixed effects, and family id as a random effect. We conducted a 100,000-permutation test to evaluate statistical significance of the relationship between the MRS and MetS in HyperGEN. In GOLDN we used a logistic mixed model to test the association between the GOLDN standardized MRS and MetS adjusting for age, sex, study site, and methylation PCs as fixed effects, and family id as a random effect. Finally, to evaluate model fit we compared the Akaike information criterion (AIC) and the Bayesian information criterion (BIC) between a basic model (age, sex, center and random effect of family) and the basic model plus the MRS within GOLDN. All statistical tests were conducted in R [12].

## Results

### Study Population Characteristics

Demographic characteristics of the HyperGEN (N=614) and GOLDN (N=995) populations - with and without MetS - are presented in Table 2. The majority of participants were female in both HyperGEN (66.61%) and GOLDN (52.26%), with an overall mean age of 49 and 50 years, respectively. Participants with MetS (MetS+) were older compared to those without (MetS-) (HyperGEN: 52 ± 10 years and 46 ±11 years, and GOLDN: 56 ±13 and 44 ±16, respectively) and more likely to be male in GOLDN and female in HyperGEN, respectively. In HyperGEN 56.2% of participants with MetS had 3 of the 5 (i.e. WC, BP, TG, HDL and FG) possible components (N=149) and fewer had 4 (N=77, 28.7%) and 5 (N=42, 15.7%) components, while in GOLDN, 47.5% of individuals met 3 out of 5 components (N=187), 35.1% had 4 of 5 (N=138) and 17.3% had 5 of 5 (N=68) components.

Table 3 shows the 6 candidate CpG association results for MetS in HyperGEN and GOLDN. With the exception of cg02650017 in *PHOSPHO1*, which was not significant in either study, the direction of association of the CpG with MetS was consistent between GOLDN and HyperGEN with at least marginal significance. Only cg18181703 in *SOC3S* was not associated with MetS in GOLDN. Both *CPT1A* cg00574958 and *ABCG1* cg06500161 were strongly associated with MetS in both studies (P<0.0001). Finally, the direction of association for *CPT1A, ABCG1, SOCS3, TXNIP* and *SREBF1* was consistent with that reported in the literature for MetS and/or related traits (Table 1).

**Table 3.** Single CPG, association results for GOLDN and HyperGEN.

|  | **Genes** | **CpG site** | **Estimate** | **SE** | **Z-score** | **P-value** |
| --- | --- | --- | --- | --- | --- | --- |
| HyperGEN*<br>Participants<br>(N=614) | *CPT1A* | cg00574958 | -0.495 | 0.113 | -4.396 | 1.10E-05 |
|  | *PHOSPHO1* | cg02650017 | -0.133 | 0.101 | -1.312 | 0.189 |
|  | *ABCG1* | cg06500161 | 0.550 | 0.113 | 4.865 | 1.15E-06 |
|  | *SREBF1* | cg11024682 | 0.570 | 0.119 | 4.777 | 1.78E-06 |
|  | *SOCS3* | cg18181703 | -0.204 | 0.095 | -2.153 | 0.031 |
|  | *TXNIP* | cg19693031 | -0.402 | 0.102 | -3.939 | 8.18E-05 |
| GOLDN**<br>Participants<br>(N=994) | *CPT1A* | cg00574958 | -0.852 | 0.113 | -7.539 | 4.72E-14 |
|  | *PHOSPHO1* | cg02650017 | 0.062 | 0.097 | 0.632 | 0.527 |
|  | *ABCG1* | cg06500161 | 0.394 | 0.099 | 3.966 | 7.31E-05 |
|  | *SREBF1* | cg11024682 | 0.270 | 0.119 | 2.269 | 0.023 |
|  | *SOCS3* | cg18181703 | -0.059 | 0.095 | -0.623 | 0.533 |
|  | *TXNIP* | cg19693031 | -0.267 | 0.092 | -2.915 | 0.004 |

\*HyperGEN model adjusted for age, sex, study site, 4 ancestry principle components, estimated blood cell counts (CD8T cells, CD4T cells, Natural Killer cells, B-cells, Monocyte cells) as fixed effects, and family id as a random effect
\*\*GOLDN model adjusted for age, sex, study site, 4 methylation principle components, and family id as a random effect.

### Risk Score Discovery and Validation

Figure 1 shows the normal distribution of the standardized MRS in the GOLDN cohort. Results from association analyses of the MRS with MetS after adjustment for covariates in HyperGEN and GOLDN are presented in Table 4. In HyperGEN, the MRS was significantly associated with MetS (permutation test p<0.0001), with each standard deviation (SD) increase in the score associated with 2.25 higher odds of having MetS (OR= 2.25; 95% CI: 1.79 -2.86). The MetS and MRS relationship was also significant in GOLDN where similarly, a 1 SD increase in the score was associated with 2.45 higher odds of having MetS (OR= 2.45; 95% CI: 2.02 -3.00). Lastly, we tested the fitness of a basic MetS model (Model 1: MetS = Age+Sex+Center+random family effect) and the basic model plus the MRS score (Model 2: MetS = Age+MRS+Sex+Center+random family effect) in GOLDN. Between Model 1 and Model 2 both AIC and BIC calculations indicate Model 2 as the best model (Table 5).

**Figure 1.**
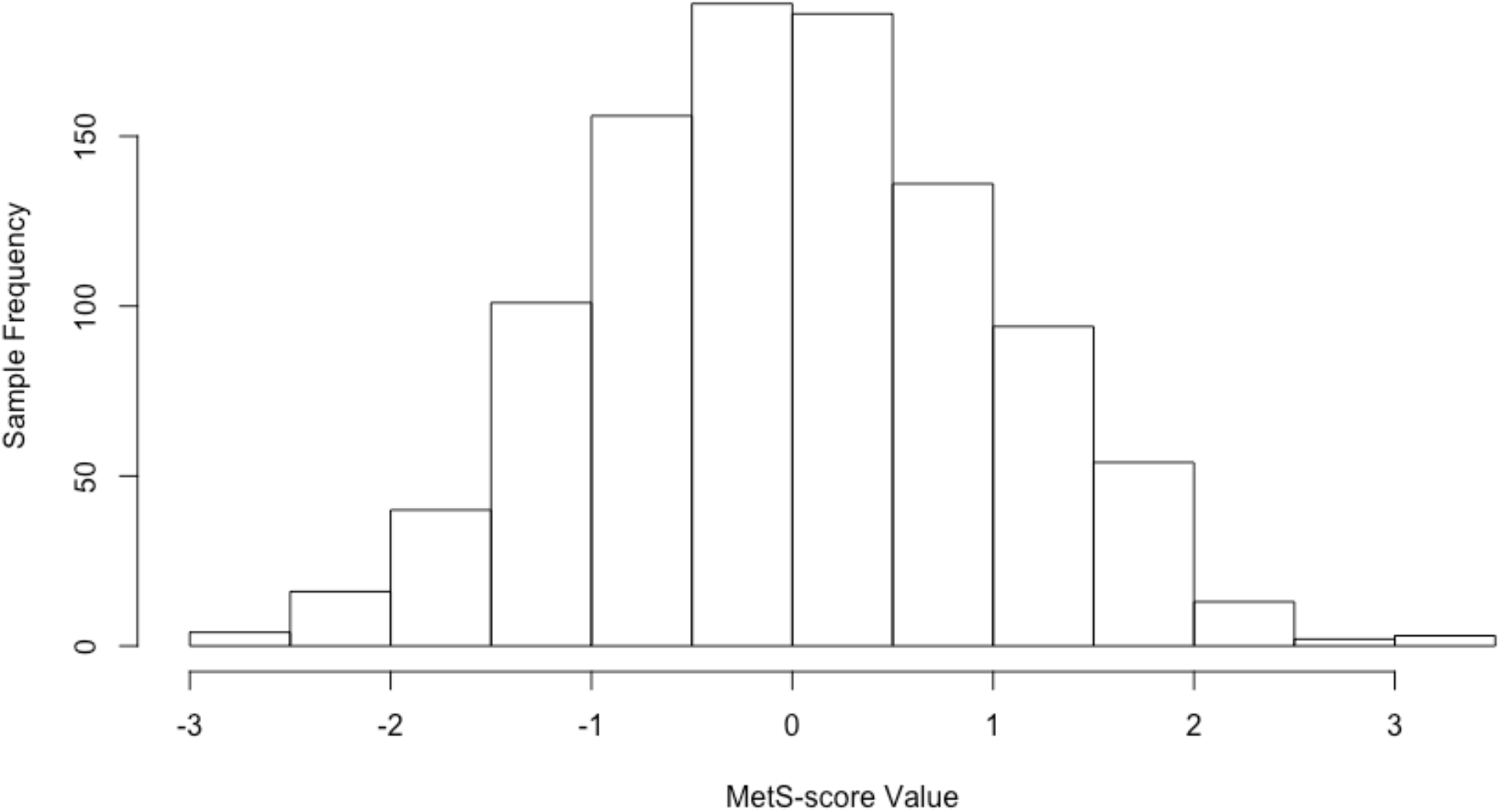
GOLDN MetS-score Distribution, N=994.

**Table 4.** Cohort-specific Association of the Methylation Risk Score (MRS) with MetS.

| **MRS** | **OR** | **P** | **95% CI** |
| --- | --- | --- | --- |
| HyperGEN* | 2.254 | 6.36E-10 | 1.80 - 2.86 |
| GOLDN** | 2.458 | <2.0E-16 | 2.03 - 3.0 |

\*HyperGEN model adjusted for age, sex, study site, 4 ancestry principle components, estimated blood cell counts (CD8T cells, CD4T cells, Natural Killer cells, B-cells, Monocyte cells) as fixed effects, and family id as a random effect.
\*\*GOLDN model adjusted for age, sex, study site, 4 methylation principle components, and family id as a random effect.

**Table 5.** GOLDN MetS and MetS-scAIC & BIC from Model 1 & 2.

|  | **AIC** | **BIC** |
| --- | --- | --- |
| **Model 1<sup>a</sup>** | 1173.99 | 1198.50 |
| **Model 2<sup>b</sup>** | 1112.61 | 1142.02 |

a) Model 1: MetS = Age + Sex + Center + Family (random effect)
b) Model 2: MetS = Methylation Risk Score + Age + Sex + Center + Family (random effect)

## Discussion

Signatures of DNA methylation associated with cardiometabolic diseases have not been widely tested for their utility in generating genomic risk scores. With better replication results across external groups and even by race these CpGs may prove useful for evaluating disease risk. Here, we introduce a six-CpG methylation risk scrore estimate for MetS that was consistent in two independent populations (HyperGEN and GOLDN). Overall, the successful performance of this MRS in two different racial populations, provides promise for future exploration of MRSs for complex disease prediction.

A substantial number of studies support the role of DNA methylation in MetS and its components. However, unlike studies of single nucleotide polymorphisms (SNPs) that have extensively considered the utility of GRS and PRS (noting many limitations, especially with respect to race), relatively few publications have included MRS [17, 18]. For instance, Hamilton *et al*, reported a positive association between an epigenetic BMI risk score and higher BMI (R^2^=0.1) in the Lothian Birth Cohort [19]. MRS for arterial stiffness measurements have been reported using data from the REGICOR and Framingham studies. In that study, two different MRS (based on alternate analytical approaches) were directly associated with arterial distensibility coefficient and inversely with pulse wave velocity [20]. Braun and others constructed a MRS in the Rotterdam study for HDL and triglycerides, finding that HDL-C levels decreased as quartiles of MRS increase, while triglyceride levels increased from the first quartile to the second quartile but remained similar for the third quartile and the fourth quartile of the MRS [21]. In another EWAS for BMI, a MRS constructed from the findings predicted future development of type 2 diabetes [22]. Finally, in a study of type 2 diabetes (∼2000 Asian Indians and ∼1000 Europeans) a score created from methylation markers at five loci (*ABCG1, PHOSPHO1, SOCS3, SREBF1*, and *TXNIP*) was associated with developing type 2 diabetes (RR of type 2 diabetes incidence 1·41 per 1 SD change in methylation score; p=1·3 × 10^−26^) [23]. Along with our study these findings suggest promise in the use of methylation scores for metabolic disease risk prediction.

MetS is strongly associated with risk for developing future diabetes and atherosclerotic and nonatherosclerotic cardiovascular disease (CVD). Though beyond the scope of this cross-sectional work, CpG sites associated with MetS may be the cause or consequence of the condition (or features of the condition). Therefore, a MetS MRS score may capture information about cumulative exposure to risk trait features. Importantly, this information could improve upon existing risk algorithms (constructed from clinical, demographic and lifestyle factors) used to predict future cardiometabolic disease. For instance, in a study set in the Bogalusa Heart Study five well-documented diabetes risk scores (non-genomic) were tested, and all showed significant associations with development of incident diabetes. These five unique risk scores differed slightly by make-up of 5–10 traditional risk factors (e.g. hypertension, smoking, family history of diabetes, age, and waist circumference), but, in general, showed good specificity but poor sensitivity. Because of the low sensitivity, the authors concluded that an opportunity remains to develop a new, more sensitive diabetes prediction tools for black and white young adults [24]. The field is similar with respect to CVD risk prediction where an excess of models and different recommendations limit algorithm use [25, 26]. Given the importance of MetS to the cardiometabolic disease landscape, and that MRS may help refine risk metrics in diverse populations for important clinical sequelae, further evaluation of these scores should be considered for disease prediction.

While there are limitations to basing a MRS for MetS from blood-based DNA methylation (due to the proxy nature of blood as a surrogate tissue for organs involved in MetS) [27, 28], the utility of blood-based DNA methylation has been proven to be highly feasible and replicable for population studies of glycemic, lipid, and other metabolic traits (Table 1). The cross-sectional nature of this study, and lack of gold standard definition for MetS are potential limitations that should be considered in future MRS assessments. However, this study strengthened by the availability of CpGs paired metabolic data in two well characterized populations enabling both discovery and validation.

In summary, we developeda MRS for MetS using existing EWAS data from two population studies of different race groups. Addition of the calculated MRS variable to a basic model of MetS further improved model fit in the study used for score validation. Given the strength of association observed in the current study and the strong body of literature surrounding the CpG loci contributing to the methylation risk score, future studies should further consider the usefulness of this metric for evaluating risk of metabolic syndrome.

## Acknowledgements

The work in this study was funded by NHLBI R01HL104135 and 15GPSPG23890000. Dr. Hidalgo is funded by 5K01HL13060904. Ms. Minniefield is funded by 3R01HL123782-02S1.

## Contributions

BH and MRI conceived of idea, contributed to writing, and were involved in oversight of analysis. BM led the statistical analysis and was involved in writing and editing. AP, MB, and HKT consulted and/or aided in statistical analysis. BH, MRI, BM, and RT conducted literature review. DKA is PI of GOLDN and HyperGEN and contributed to editing/writing.

## Data sharing

Genomic data obtained for this study can be found on dbGap or by special request to the respective study principal investigators.

## Notes

### Competing Interest Statement

The authors have declared no competing interest.

